# Modulation of cortical slow oscillatory rhythm by GABA_B_ receptors: an experimental and computational study

**DOI:** 10.1101/2019.12.14.866442

**Authors:** M Perez-Zabalza, R Reig, J Manrique, D Jercog, M Winograd, N Parga, MV Sanchez-Vives

## Abstract

Slow wave oscillations (SWO) dominate cortical activity during deep sleep, anesthesia and in some brain lesions. SWO consist of Up states or periods of activity interspersed with Down states or periods of silence. The rhythmicity expressed during SWO integrates neuronal and connectivity properties of the network and it is often altered in neurological pathological conditions. Different mechanisms have been proposed to drive the transitions between Up and Down states, in particular, adaptation mechanisms have been proposed to contribute to the Up-to-Down transition. Synaptic inhibition, and specially GABA_B_ receptors, have also been proposed to have a role in the termination of Up states. The interplay between these two potential mechanisms, adaptation and inhibition, is not well understood and the role of slow inhibition is not yet clear regarding the full cycle of the slow oscillatory rhythm. Here we contribute to its understanding by combining experimental and computational techniques. GABA_B_ receptors-blockade not only elongated Up states, but also affected the subsequent Down states, and thus the whole cycle of the oscillations. Furthermore, while adaptation tends to yield a rather regular behavior, GABA_B_ receptors-blockade decreased the variability of the sequence of Up and Down states. Interestingly, variability changes could be accomplished in two different ways: either accompanied by a shortening or by a lengthening of the duration of the Down state. Even when the most common observation is the lengthening of the Down states, both changes are expressed experimentally and also in numerical simulations. Our simulations suggest that the sluggishness of GABA_B_ receptors to follow the excitatory fluctuations of the cortical network can explain these different network dynamics modulated by GABA_B_ receptors.

## Introduction

Different brain states are characterized by diverse patterns of spontaneous activity. The interplay between neuromodulators, receptors, intrinsic properties, and connectivity helps explain neuronal discharge, and as a network property, the emergence of different frequencies of oscillatory activity. During slow wave sleep, the synchronized discharge of a large number of cortical neurons explains the increase in voltage fluctuations observed during EEG recordings, whereas wakefulness and REM sleep can be identified by lower-voltage amplitudes (Aserinsky and Kleitman, 1953), reflecting a decrease in the number of cortical neurons discharging simultaneously. We also know that the degree of network synchronization/ desynchronization changes between different brain states and is precisely modulated during wakefulness as well (Poulet and Petersen, 2008; Okun et al., 2010; Harris and Thiele, 2011).

The modulation of network synchronization has been related to different brain functions. During slow wave oscillations (SWO) high levels of synchronization are involved in synaptic and cellular homeostasis, as well as memory formation (Hoffman et al., 2007; Diekelmann and Born, 2010; Tononi and Cirelli, 2014). It has been suggested that, during wakefulness, synchronization facilitates the transfer of information between distal neurons, providing temporal coordination for specific neuronal assemblies (Varela et al., 2001; Doesburg et al., 2010; Tononi and Cirelli, 2014). On the other hand, low levels of synchronization are observed during alert or attentional states, short-term memory tasks, or movements, at local cortical areas (Stancák and Pfurtscheller, 1996; Klimesch et al., 2007; Okun and Lampl, 2008; Doesburg et al., 2010). Malfunctions controlling neural synchronization at different oscillatory frequencies are related to several neurological diseases such as Alzheimer’s (Busche et al., 2015; Castano-Prat et al., 2019), early aging (Castano-Prat et al., 2017), Parkinson’s (Little and Brown, 2014), autism (Rubenstein and Merzenich, 2003), Williams Beuren syndrome (Dasilva et al., 2019), or Down Syndrome (Ruiz-Mejias et al., 2016), among others. Although the control of the degree of synchronization of neural activity is essential in order to understand normal and pathological brain function, there are still open questions regarding the basic mechanisms underlying network synchronization.

SWO (<1Hz) are organized into alternating periods of activity and silence–Up and Down states, respectively (Steriade et al., 1993). This spontaneous neural activity can be recorded from both isolated cortical slabs *in vivo* (Timofeev et al., 2000) and in cortical brain slices *in vitro* (Sanchez-Vives and McCormick, 2000) implying that the cortical network can generate SWO on their own (i.e., without external input). Indeed, SWO are also expressed in clinical conditions where a “cortical island” occurs as a result of a lesion (Gloor et al., 1977) or perilesional around acute ischemic cortical stroke, where SWO can persist for months or even years (Butz et al., 2004). This capability of the disconnected cortical network to generate highly similar SWO independently from the size of the cortex involved, has led to the suggestion that this SWO is the default emergent activity pattern from the cortical network (Sanchez-Vives and Mattia, 2014; Sanchez-Vives et al., 2017). Cortical SWO either in sleep or in deep anesthesia are characterized by a high degree of network synchronization, where large populations of neurons are engaged, shaping the slow wave sleep cycle (Bullock and McClune, 1989; Steriade et al., 1993; Destexhe et al., 1999; Ruiz-Mejias et al., 2011; Bettinardi et al., 2015). This synchronization can be explained by a combination of excitatory and inhibitory input that cortical neurons receive during Up states, which results in the depolarization of neuron membrane potential that, in turn, often generates bursts of action potentials. On the other hand, cortical activity during Down states remains rather silent. Such patterns of active and silent cortical activity depend on the balance between recurrent excitation and local inhibition (Sanchez-Vives and McCormick, 2000; Compte et al., 2003, 2009; Shu et al., 2003). However, the precise biophysical mechanisms underlying the inhibitory modulation of SWO is not fully understood.

In order to understand spontaneous SWO, several studies have proposed potential mechanisms responsible for their generation (transition from Down to Up states), maintenance, and termination (transition from Up to Down states). In terms of finalization, four main different mechanisms have been proposed that mediate the transition from Up to Down states: firing rate adaptation (Compte et al., 2003; Sanchez-Vives et al., 2010), short-term synaptic dynamics (Timofeev et al., 2000; Melamed et al., 2008; Benita et al., 2012), ATP-dependent homeostatic mechanisms mediated by ATP-modulated potassium (K_ATP_) channels (Cunningham et al., 2006), and pre- and post-synaptic GABA_B_ (gamma-aminobutyric acid B) receptor activation (Parga and Abbott, 2007; Mann et al., 2009; Wang et al., 2010; Craig et al., 2013) (Sanchez-Vives et al., in review). It is plausible that more than one of these mechanisms interact and contribute to the termination of the Up-to-Down state transition; however, this is not yet well understood due to a paucity of experimental and modeling work addressing this issue. Although there are some indications that firing rate adaptation is at least partly responsible for the termination of Down states (i.e. generation of SWO) (Sanchez-Vives et al., 2010), it is not known to which extent this is the dominant mechanism. Interestingly, modeling work has shown that GABA_B_ dynamics have the correct timescale to contribute to the Down state transition (Parga and Abbott, 2007) and that they could interact with firing rate adaptation to modulate the termination of Up states.

Here we combined biological and computational experiments to elucidate which mechanisms underlie modulation of SWO. More specifically, we empirically studied the role of GABA_B_ receptors in controlling the Up to Down state transitions, Up and Down states durations and variability, and their impact on the global synchronization of spontaneous activity. We also modeled and simulated two different network behaviors and tested hypotheses in order to understand the role of each of the possible biophysical mechanisms in the modulation of SWO; finally, we tested the predictions obtained from the experimental observations in our models.

## Methods

### Experimental procedures

To empirically study the role of GABA_B_ receptors in the modulation of SWO, *in vitro* experiments were performed on 37 ferret cortical slices (4- to 10-month-old, either sex). Ferrets were deeply anesthetized with sodium pentobarbital (40 mg/kg) and decapitated. Their brains were quickly removed and placed in ice-cold cutting solution (4–10°C). Coronal slices (thickness: 400 µm) of the primary visual cortex (*n*=24) and prefrontal cortex (*n*=13) were cut on a vibratome.

A modification of the sucrose-substitution technique developed by (Aghajanian and Rasmussen, 1989) was used to increase tissue viability, as in (Sanchez-Vives and McCormick, 2000). After preparation, slices were placed in an interface-style recording chamber (Fine Science Tools, Foster City, CA) and bathed in artificial cerebrospinal fluid (ACSF) containing (in mM): NaCl, 124; KCl, 2.5; MgSO4, 2; NaHPO_4_, 1.25; CaCl_2_, 2; NaHCO_3_, 26; and dextrose, 10; and was aerated with 95% O_2_ and 5% CO_2_ to a final pH of 7.4. Bath temperature was maintained at 34–35°C. After 2 h recovery, the ACSF was replaced by “*in vivo*-like” ACSF (Sanchez-Vives and McCormick, 2000) containing (in mM): KCl, 3.5; MgSO_4_, 1; CaCl_2_, 1; the remaining components were the same as those just described. Extracellular, unfiltered recordings were obtained by means of tungsten electrodes through a Neurolog system (Digitimer) amplifier. Intra and extracellular recordings were digitized, acquired using a data acquisition interface (CED) and software (Spike2) from Cambridge Electronic Design.

To study the effects of GABA_B_ blockade on SWO, GABA_B_ antagonist CGP 35348 (200µM) (Tocris) was added to the bath. Baseline recordings before CGP 35348 application were used as the control conditions.

In a subset of experiments (*n*=13), we recorded extracellularly from deep cortical layers before and after eliminating layer 1 from the cortical circuit. To isolate the circuit from long-lasting connections, a cut was made between layers 1 and 2/3, parallel to the white matter, and two additional cuts were made perpendicular to the white matter (Fig. S1). To study GABA_B_ blockade before and after isolating the circuit, GABA_B_ blocker CGP 35348 (200µM) (Tocris) was added to the bath (*n*=11). Baseline recordings before cutting and before CGP 35348 application were used as the control conditions.

### Spike recording and analysis

Extracellular multiunit recordings were obtained with 2–4 MΩ tungsten electrodes (FHC, Bowdoinham, ME). Multiunit activity (MUA) was estimated as the power change in the Fourier components at high frequencies of the recorded local field potentials (LFP). High-frequency components of LFP can be seen as a linear transform of the instantaneous firing rate of the neurons surrounding the electrode tip. We then assume that the normalized LFP spectra provides a good estimate of the population firing rate, given that Fourier components at high frequencies have power densities proportional to the spiking activity of the involved neurons.

Several parameters were quantified from the transformed signal (log(MUA)): (1) the maximum firing rate in the Up state was the peak of the average log(MUA) in the time interval (−0.5, 2.5 ms) around the Up state onset; (2) the upward and downward transition slopes were the gradients of the linear fits of the average log(MUA) in the time intervals (−10, 25 ms) and (−25, 10 ms), respectively, around the Up-state onset and offset, respectively; and (3) to measure the variability of SWO, we calculated the coefficients of variation (CV) (standard deviation/mean) of Up state duration, Down state duration and oscillation cycle duration (Up state duration + Down state duration). Data are reported as mean±SD. All off-line analyses were implemented in MATLAB (The MathWorks Inc., Natick, MA).

### Model and theoretical procedures

To better understand the role of GABA_B_ receptors in the modulation of SWO, we simulated networks of integrate-and-fire neurons, with the addition of a nonlinear membrane current, receiving synaptic input composed of slow and fast excitatory and inhibitory conductances (Parga and Abbott, 2007). These simulated networks consist of random connections with finite range. Each neuron is described by its membrane potential *V* which, below its threshold value, evolves according to the equation

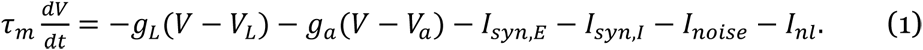

Here, *τ_m_* is the membrane time constant, *g_L_* is the leak conductance and *V_L_* is the leak reversal potential. The adaptation current, *g_a_* (*V* − *V_a_*), only affects the excitatory neurons. Its conductance, *g_a_*, decays exponentially with a time constant, *τ_a_*, until a spike is fired. When this happens, the adaptation conductance is augmented by an amount *g_a_*. *I_syn,E_* and *I_syn,I_* are the excitatory and inhibitory synaptic currents, respectively. *I_noise_* is an external noise current. *I_nl_* describes a nonlinear property of the neuron (see below). A neuron fires whenever its membrane potential *V*(*t*) reaches the spike generation threshold *V_th_*. At this point, an action potential is triggered, and the potential *V*(*t*) is reset and kept at a value *V_reset_* during a refractory period *τ_ref_*. Two excitatory (AMPA, NMDA) and two inhibitory (GABA_A_, GABA_B_) synaptic currents are included as *I_syn,E_(t) = g_AMPA_(V (t) − V_AMPA_) + g_NMDA_(V (t) − V_NMDA_)*

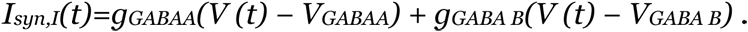

When a neuron fires an action potential, the synaptic conductances of its postsynaptic neurons are modified by an amount *Δg_X_* (*X*=AMPA, NMDA, and GABA_A_, GABA_B_). Otherwise, the synaptic conductances decay exponentially, with synaptic time constant *τ_X_*. Nonlinearities characterizing NMDA and GABA_B_ receptors are not considered; the emphasis in this model is on the time scales of the conductances. The nonlinear membrane current is a simple way of accounting for the neuron’s intrinsic properties. It is described as

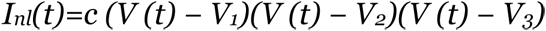

where *V*_1_< *V*_2_< *V*_3_ and *c* determines the strength of the current. In the absence of noise, *I_nl_* induces three fixed points, one of them being unstable. Fluctuations produced by the noise term and by the synaptic currents allow the neuron’s membrane potential to alternate in a bistable fashion instead of being stuck at stable fixed-point values. Each neuron receives independent noise currents *I_noise_* consisting of two filtered Poisson trains, one excitatory and one inhibitory. This current is parameterized by two unitary conductances (*Δg_syn,E_* and *Δg_syn,I_*), two Poisson rates and the time constants of the filter (*τ_NMDA_* and *τ_GABAB_*).

### Parameter values

We simulated two networks (which we call the typical and atypical networks) in order to reproduce the experimentally observed data. These networks differ in parameters related to synaptic and adaptation properties (see below).

We also simulated networks with parameters as those of the typical network but with different values of the characteristic time of the adaptation current. Networks contained 4000 neurons of which 17% were inhibitory and the rest, excitatory. Pairs of neurons separated by a distance shorter than a certain radius were connected with a probability of 2%. This radius was chosen in such a way that, on average, each neuron was connected to 25 other neurons. The network size was 50 × 80 neurons, with periodic boundary conditions. All the neurons had a membrane time constant *τ_m_* = 20 ms and a refractory time *τ_ref_* = 5 ms. Other passive properties were distributed uniformly. The membrane threshold *V_th_* took values of −45±2 mV, the reset potential *V_reset_* of −55±1 mV, and the leak potential *V_L_* of −68±1 mV. The parameters of the nonlinear current were *c*=0.03 mV^−2^, *V_1_*=−72±2 mV, *V_2_=*−58±2 mV and *V3*=−44±2 mV. AMPA and NMDA currents were present in all excitatory synapses. Similarly, we assigned GABA_A_ receptors to 100% of the inhibitory synapses but GABA_B_ receptors to only 70% of them.

The parameters of the noise model were: *Δg_syn,E_*=0.09, *Δg_syn,I_*=0.18 for the conductances and *υ_syn,E_*=66.66 Hz, *υ_syn,I_*=24.31 Hz for the frequency rates. The synaptic time constants were *τ_AMPA_*=2 ms, *τ_NMDA_*= 100 ms, *τ_GABAA_* =10 ms, and *τ_GABAB_*=200 ms. All conductances are measured in units of the excitatory leak conductance (which we took as *g_E,L_*=10 nS). *Δg_E,AMPA_=*0.54, *Δg_E,NMDA_*=0.04, *Δg_E,GABAA_*=1.00, *Δg_E,GABAB_*=0.18. For inhibitory neurons, *Δg_I,AMPA_*=0.38 (typical network) or *Δg_I,AMPA_*=0.57 (atypical network), *Δg_I,NMDA_*=0.04, *Δg_I,GABAA_*=0.02, *Δg_I,GABAB_*=0.017 and *g_I,L_*=1.4. In addition, for excitatory neurons, *Δg_a_*=0.03, *V_a_=*−80 mV and *τ_a_*= 1900 ms (typical network) or *τ_a_*=2500 ms (atypical network). The reversal potentials for the inhibition, *V_GABAB_* and *V_GABAA_* fall uniformly within the values −90±2 mV and −80±2 mV, respectively. *V_AMPA_* and *V_NMDA_* are both set to zero. We also considered another example of typical network with the same parameters as the one described above except for Δg_E,NMDA_=0.05, Δg_I,NMDA_ =0.05 and *τ_a_*= 4000 ms.

### Up and Down transition detection algorithm

This algorithm provided criteria for determining when the network moved from one state to another. The criteria can be summarized as follows: (1) Up-to-Down transitions: at a given time, the number of spikes of each neuron in a window of 60 ms was measured. If every cell fired less than two spikes, the transition to the Down state took place. (2) Down-to-Up transitions: if the percentage of neurons that fired in windows of 60 and 100 ms was at least 10% and 30%, respectively, then the transition to the Up state occurred (Luczak et al., 2007).

### Correlation functions and coefficients of variation (CV)

We calculated spike correlograms as the average over a subpopulation of 200 randomly selected neurons of the pair-wise correlation function

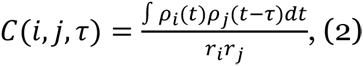

where *ρ_i_*_(*j*)_(*t*) and *r_i_*_(*j*)_ are the spike train and the firing rate, respectively, of neuron *i* (*j*), and *τ* is the time lag. Correlation functions of the currents were computed as:

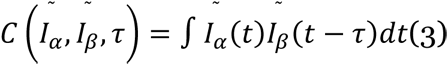

where 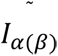 is the population average of the current *I*_α(*β*)_. Both *C*(*i*, *j*, *τ*) and 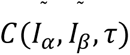 were normalized to their value at their respective peaks. CVs of the duration of the Up states, the Down states and the cycle were defined as the ratio between the standard deviation and the mean (standard deviation/mean).

### Simulation

Simulation times were typically 1200 s. Differential equations were integrated using the Euler method with an integration step of *Δt*=0.1 ms. To obtain Fig. 7A, we ran eight simulations of 150 s for each value of the parameter *τ_a_*, with different realizations of the noise. Statistical errors in these graphs were computed as the SD of the values obtained in each simulation. All codes were written in C and run under the Linux operating system.

## Results

The cortical network *in vitro* preserves the mechanisms to generate spontaneous rhythmic neural activity, namely SWO, organized into Up states (active periods), and Down states (silent periods). Recordings from ferret cortical slices revealed spontaneous SWO (Fig. 1A).

**Figure 1.**
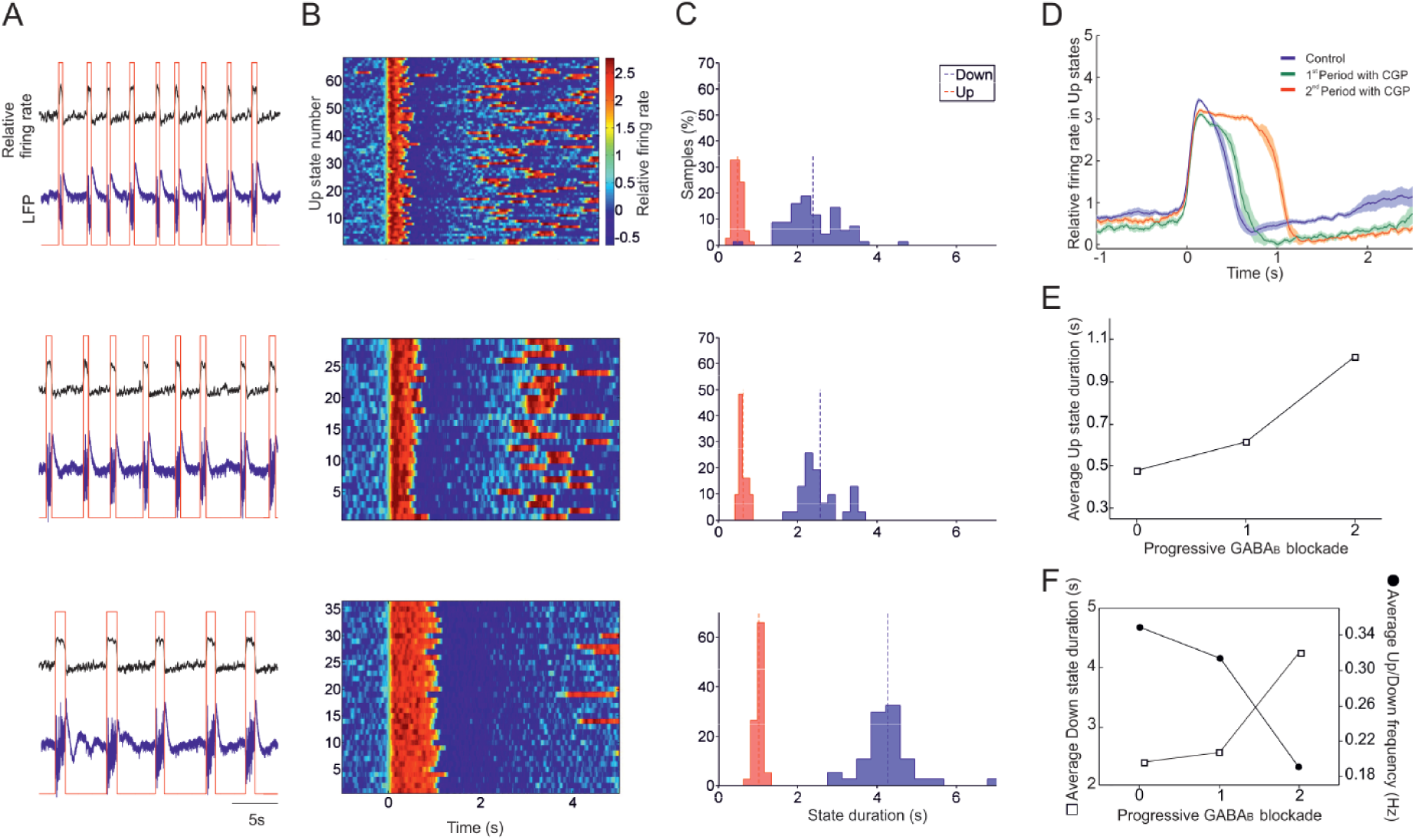
Effects of progressive inhibition blockade on slow oscillations. Typical network. **A.** Raw signal (blue trace) and relative firing rate (black trace; see Methods). Up states were detected from the relative firing rate (red trace). From top to bottom: control activity and two consecutive periods after application of 200 µM CGP 35348. Time scale is the same for all panels in A. **B.** Raster plots of the relative firing rate are represented for control activity and for 200 µM CGP 35348 corresponding to the ones in A. The firing rate is color coded. Time scale is the same for all panels in B. **C.** Histograms of the Up and Down duration for control activity and 200 µM CGP 35348 corresponding to the ones in A. **D.** Average relative firing rate for Up states during the control and 200 µM CGP 35348 corresponding to the ones in A. The shadow corresponds to the s.e.m. **E.** Up state duration increases with CGP 35348. **F.** Down state duration increases whereas the oscillation frequency decreases. Empty squares are average Down state duration and filled circles are average Up/Down frequency.

### Effects of GABA_B_ blockade on the Up / Down state cycles

The baseline frequency of the slow oscillations was 0.35 ± 0.02 Hz, with a duration of Up/Down states of 0.56 ± 0.05 s and 2.55 ± 0.17 s respectively. GABA_B_ antagonist CGP35348 applied to the bath resulted in the gradual blockade of slow inhibition and induced several changes in the Up and Down states of the cortical slices (*n*=37) (see Fig. 1 for a particular slice). In agreement with previous studies (Mann et al., 2009), a prominent and consistent change in Up and Down state properties upon GABA_B_ blockade was the elongation of the Up states (Fig. 1A-F).

The Up state elongation resulting from GABA_B_ blockade was observed in 34 out of the 37 slices (Fig. 1D, 1E, Fig. 2A and 3D, 3E), while there were no changes in the remaining three cases. This elongation was on average to 182% the original Up state duration and was observed independently of the original duration in the control (baseline) condition, which ranged between 0.2 and 1.3 seconds (Fig. 2A). The elongation of Up states following GABA_B_ receptor blockade suggests that these receptors participate in the termination of the Up states, as it has been previously proposed (Parga and Abbott, 2007; Mann et al., 2009; Wang et al., 2010; Craig et al., 2013), although it could also be secondary to the alteration of the excitatory/inhibitory balance during the Up state and subsequent modulation of the firing rate during Up states (Compte et al., 2003; Mattia and Sanchez-Vives, 2012). We explore these possibilities next.

**Figure 2.**
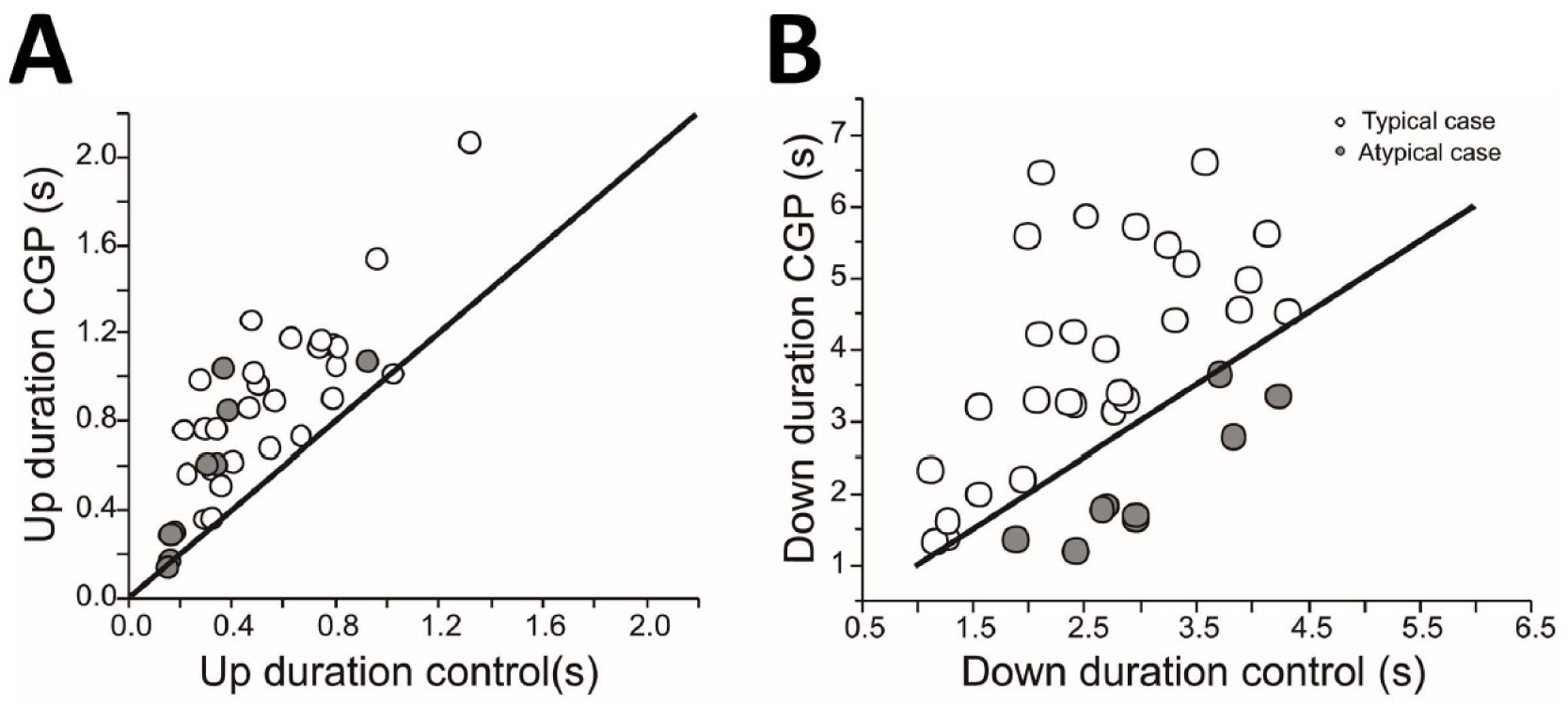
Modulation of Up and Down states by GABAB-receptors. **A**, scatter plot of the duration of Up states in control *versus* blockade of GABAB receptors with 200 µM CGP 35348. In all cases the Up states duration increases or in few cases remains the same. **B**, same for Down state duration. In the two panels, the imaginary line is the one corresponding to the absence of changes (bisecting line). We define as “typical networks” those were the Down state duration gets elongated (empty circles), and “atypical networks” those in which the Down states got shorter (gray-filled circles).

**Figure 3.**
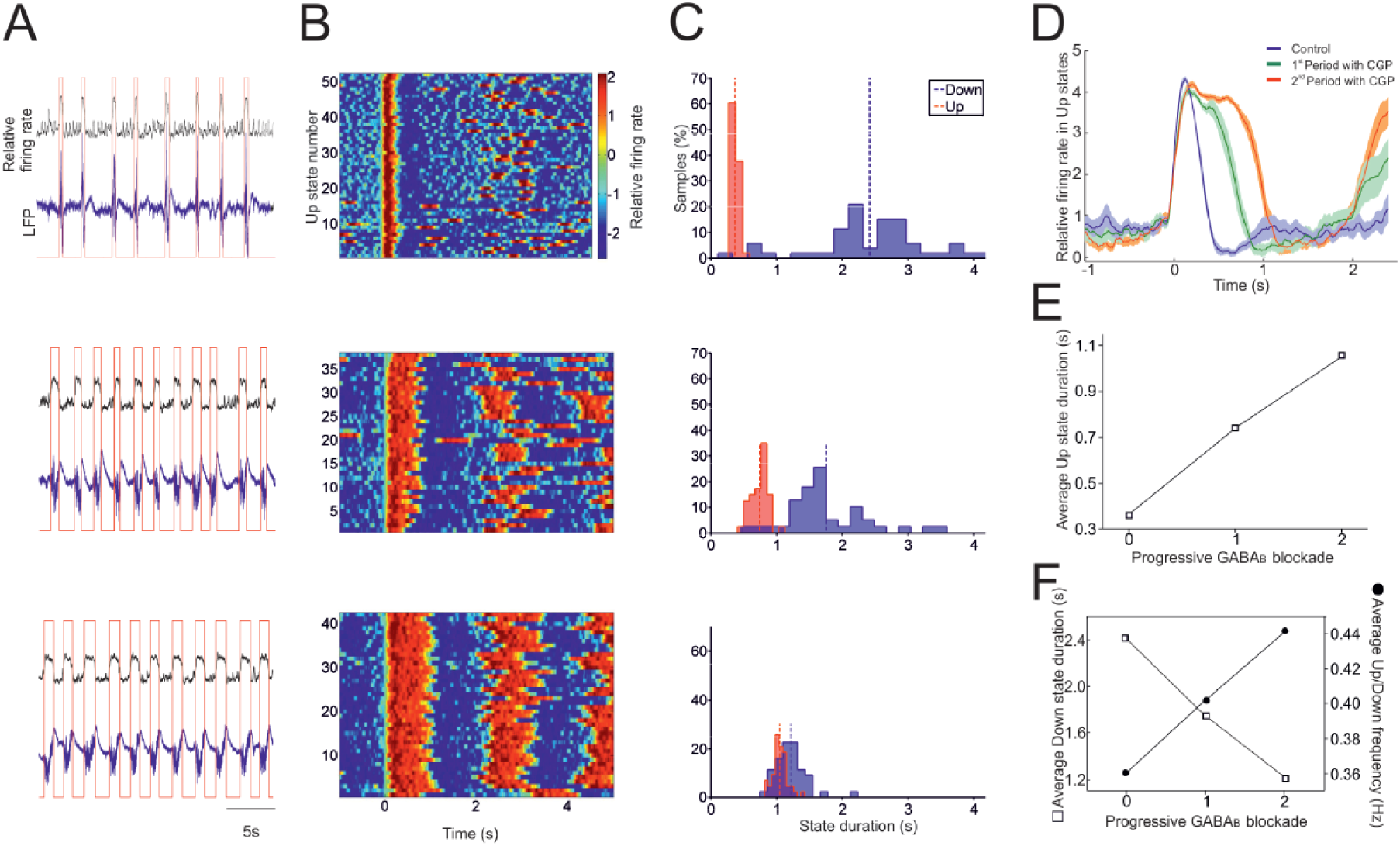
Effects of progressive inhibition blockade on slow oscillations. Atypical network. **A.** Raw signal (blue trace), relative firing rate (black trace; see Methods) and detected Up and Down states (red trace). From top to bottom: control activity and two consecutive periods after application of 200 µM CGP 35348. **B.** Raster plots of the relative firing rate are represented for control activity and for 200 µM CGP 35348 corresponding to the ones in A. The firing rate is color coded. **C.** Histograms of the Up and Down duration for control activity and 200 µM CGP 35348 corresponding to the ones in A. **D.** Average relative firing rate for Up states during the control and 200 µM CGP 35348 corresponding to the ones in A. The shadow corresponds to the s.e.m. **E.** Up state duration increases with CGP. **F.** Down state duration decreases whereas the oscillation frequency increases.

Down states were also globally elongated as a result of GABA_B_ blockade. The average elongation of the Down state (*n*=37) was to the 138% of the original value (Fig. 1, Fig. 2). However, when recordings were looked at individually, we observed two patterns (or groups): even though the most common effect was the elongation of the Down states (*n*=28, Fig. 1; Fig. 2B), in some cases Down states were indeed shortened by GABA_B_ blockade with CGP35348 (*n*=9, Fig. 2B; Fig. 3). We called the first group “typical” (Fig. 1) and the second one “atypical” (Fig. 3), and this is how we will refer to them in the rest of the study. The shortening of Down states could occur even when the Up states were elongated similarly to the typical case in Fig. 1, resulting in the overall increase of the frequency of the oscillatory cycle (Fig. 3F).

Up and Down states are dynamically related (Compte et al., 2003; Sanchez-Vives et al., 2010; Mattia and Sanchez-Vives, 2012); it is therefore intriguing that the same transformation of the Up states (elongation) is followed by two different transformations of the Down states (typical elongation, or atypical shortening) (Fig. 2A, B). A possible functional explanation of these results could be that SWO in typical and atypical networks were different to start with. Indeed, we found that in control conditions, the Up state duration was significantly shorter in atypical than in typical networks (atypical: 0.32±0.08 s; typical: 0.56±0.05 s; *p*<0.05), although this observation still lacks a mechanistic explanation. We further explore the possible dynamical mechanisms in our computer model below.

Interestingly, GABA_B_ receptor blockade strikingly increased the regularity of the SWO. This effect was obvious in the autocorrelations of the activity before and after GABA_B_ blockade, where the time constant of an exponential fitted to the peaks (coherence time (Dowse, 2009) became slower under GABA_B_ blockade (Fig. 4A,B, insets in (a)).

**Figure 4.**
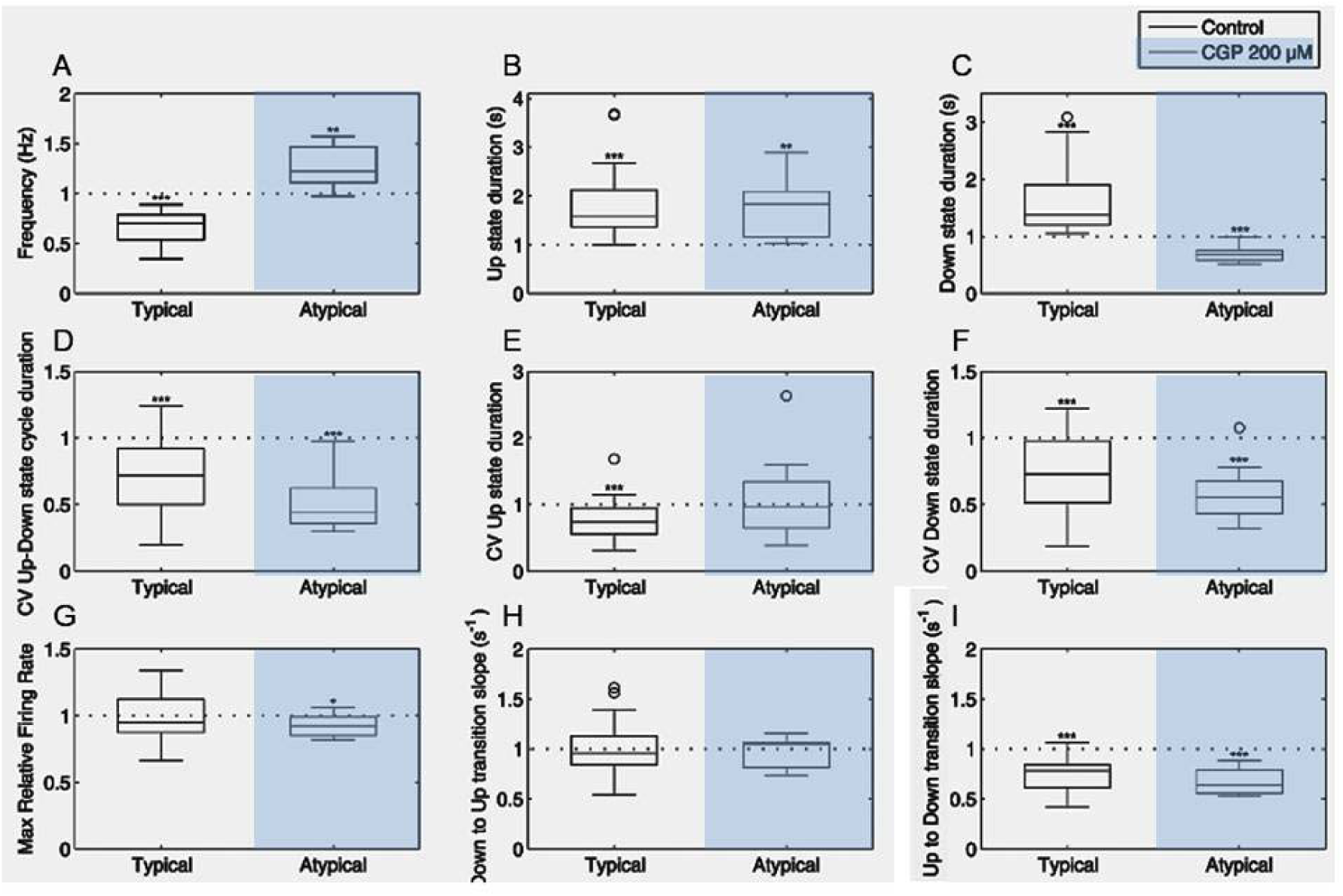
Effect of GABA_B_ blockade on the variability of slow oscillations. **A**. ‘Typical’ network (same recording as Fig. 1). (a) Autocorrelograms illustrating the transformation of the emerging activity for control activity and for two consecutive periods after application 200 μM CGP35348 (left to right). Inset: The decay envelope of the autocorrelogram is a function of the long-range regularity in the signal (Chatfield, 1980). (b) Measure of the decay envelope of the autocorrelogram. (c) Rhythmicity Index. (d) CV of Up/Down cycle, Up state duration and Down state duration. **B.** ‘Atypical’ network (same recording as Fig.2). Same parameters that (A).

The change in variability of the durations of Up and Down states and oscillatory cycles was also quantified by the coefficient of variation (CV). The CV of Up states in atypical networks, and also the CV of Down states and of the oscillatory cycle, were significantly decreased by GABA_B_ blockade in both typical (Fig. 4Ad) and atypical (Fig. 4Bd) networks. Compared with control values, the decrease of the CV for the typical case was to 85% and 68% for the Up and Down states, respectively, and to 66% for the complete oscillatory cycle (Fig. 5).

**Figure 5.**
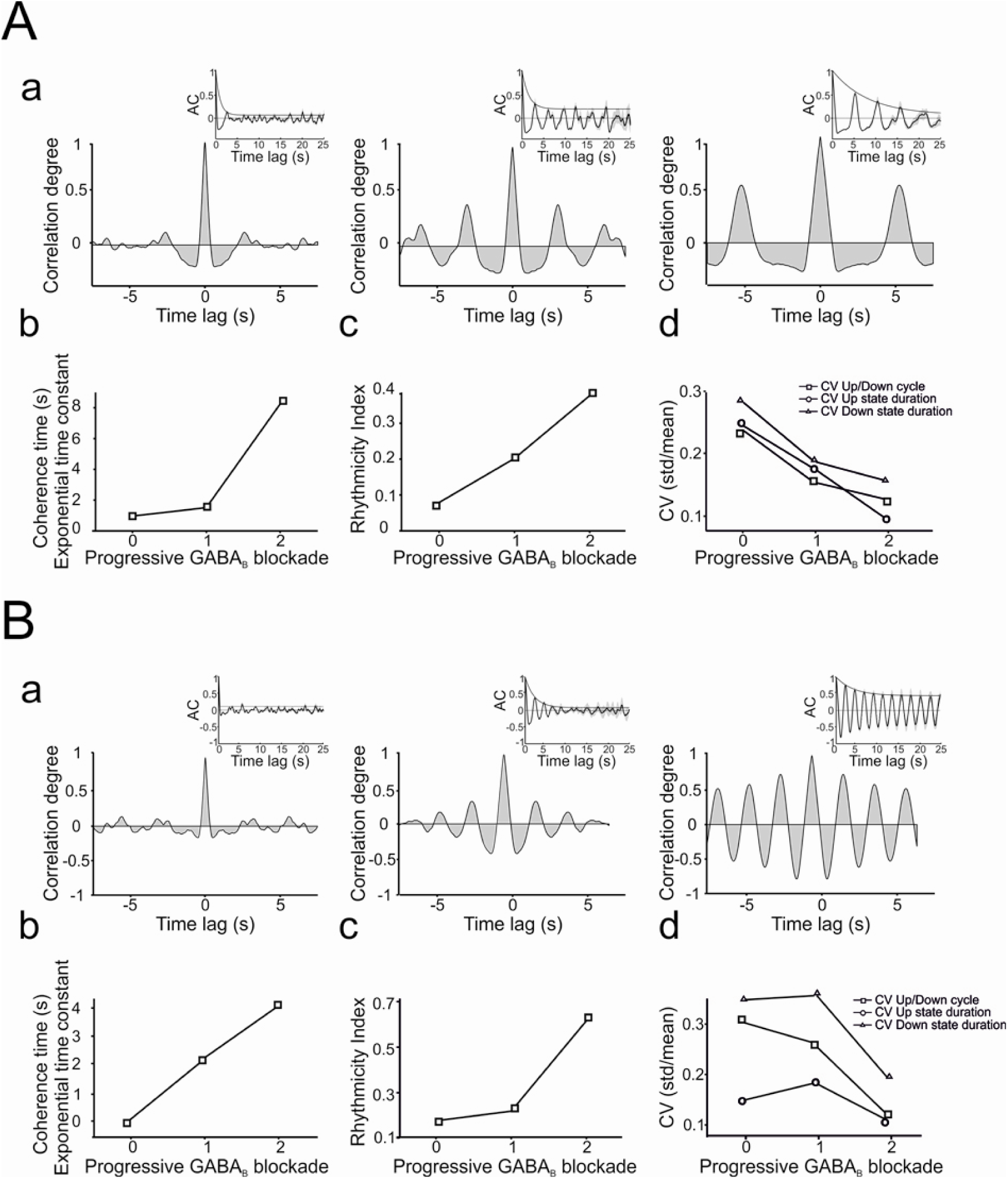
Relative changes of Up and Down states properties after removal of slow synaptic inhibition on typical (*n*=28) and atypical (*n*=9) network. Average values of different parameters calculated. A Student’s *t*-test was used to evaluate how different the normalized averages were from 1.0., *p* < 0.05 (*); *p* < 0.01 (**); *p* < 0.001 (***).

In the case of the atypical case there was a significant decrease in CV of 58% for the Down state, and 51% for the oscillatory cycle. These results indicate that the physiological activation of GABA_B_ receptors introduces variability and dynamical richness in the spontaneous SWO. Hence, activation of GABA_B_ receptors does not only contribute to the modulation of the duration of the Up states, but also affects the network dynamics by controlling the duration of the Up/Down states and by disrupting the regularity of the SWO (Fig. 5).

The firing rate during Up states is in some cases the link that explains the Up/Down state relative durations. This can be the case when the mechanisms of termination of Up states are activity-dependent, for example the activation of sodium- and calcium-dependent K^+^ currents (Compte et al., 2003). In cases where GABA_A_ receptors are blocked, the decrease in fast inhibition results in high firing rates during Up states that efficiently activate sodium- and calcium-dependent K^+^ currents that shorten the Up states and elongate Down states (Sanchez-Vives et al., 2010). We explored if this was the case when GABA_B_, and not GABA_A_, receptors were blocked. We did not find any significant difference when comparing the firing rate in Up states before and after GABA_B_ blockade (Fig. 5G; *n*=37). This finding suggests that the GABAergic control of firing rate in Up states mostly occurs through GABA_A_ receptors, while the role of GABA_B_ receptors on firing rate is negligible but noticeable on the termination of Up states and on network dynamics.

To understand the dynamics of the oscillatory activity in the cortical network is useful looking into the transitions between states, such as the slopes of the Down-to-Up and Up-to-Down transitions. The Down-to-Up transition slope, which corresponds to the recruitment of the local network (Reig et al., 2010; Sanchez-Vives et al., 2010) was not affected by GABA_B_ receptor blockade. Interestingly, the Up-to-Down (downward) transition slope was significantly decreased (Fig. 5I) both in typical and atypical networks, meaning that the finalization of the SWO cycle was slower when GABA_B_ receptors were blocked, further supporting the role of these receptors in the termination of Up states.

We investigated the possible role of cortical layer I in modulating the spontaneous Up and Down states in infragranular layers before and after GABA_B_ blockade. To this end, we recorded spontaneous SWO before and after eliminating layer 1 from the cortical network by cutting the slice between layers 1 and 2/3, with and without GABA_B_ blockade with CGP 35348 (Fig. S1A, *n*=11). GABA_B_ blockade resulted in a significant elongation of the Up states even in the absence of layer 1, similar to what occurred in control slices (Up state duration: control 0.47±0.05 s; layer 1 eliminated 0.34±0.04; layer 1 eliminated + CGP 35348 0.53±0.04, Fig. S1). This result parallels a previous study showing that, during spontaneous oscillatory activity, GABA_B_ contributes to the Up-to-Down state transitions without the influence of layer 1 (Craig et al., 2013).

In our study we also explored whether different circuits can display diverse effects after blocking GABA_B_ receptors. For this, we recorded the spontaneous SWO in supra- and infragranular layers simultaneously with and without CGP 35348. The results did not show differences between layers (*n*=16). The effects on supragranular layers were not different to the ones described for infragranular layers in Figs 1 and 2. These results show that GABA_B_ receptors strongly modulate the spontaneous neural activity in different layers and cortical areas as reported above.

In conclusion, from the experimental results we observed that the blockade of GABA_B_ receptors decreased the variability (CV) of the duration of Up and Down states as well as of the complete oscillatory cycle, suggesting that GABA_B_ receptor activation plays a key role desynchronizing network activity. We also observed a prominent and consistent elongation of Up states as a consequence of GABA_B_ receptor blockade, confirming that GABA_B_ activation participates on the termination of Up states. The fact that the slope of the Up-to-Down state transition became slower when GABA_B_ receptors were blocked is in agreement with the suggested role of these receptors in Up state termination proposed by (Mann et al., 2009). In most cases the Up state elongation after GABA_B_ blockade occurred concurrently with an elongation of the subsequent Down states (typical network), although in one-quarter of the cases the Down states shortened (atypical network). We designed a computer model of the cortical network that reproduces these observations and proposes a mechanistic explanation for them, proposing a role for GABA_B_ receptors in the dynamics of the SWO.

### The model: description of its basic properties

First, we present the basic features of the SWO generated with our model. We generated two sample networks, one responding to GABA_B_ receptor blockade in a typical and the other in an atypical way. We next explored the most remarkable effects of GABA_B_ receptors that were reported experimentally: modulation of the duration of the Up states and modulation of the regularity of the oscillations.

We generated several sample networks with fixed connectivity but differing in the precise realization of the connectivity matrix and in the value of some parameters (see Methods for details). Two examples of networks (typical and atypical) were defined such that they had approximately the same cycle duration in the control condition (Fig. 6A and 6B). For both networks, in all the generated samples and in all the slices recorded in our experiments, blocking GABA_B_ receptors did not suppress the SWO, and the duration of the Up state became longer. For the typical network (Fig. 6A), the duration of the Down states increased as it did in the typical experimental cases. For the atypical network (Fig. 6B), the duration of the Down states decreased as it occurred in the atypical experimental cases as well. Notice that for the two networks, the histogram of the duration of Down states (panels b and e in Figs. 6A and 6B) had a larger dispersion when GABA_B_ receptors were not blocked (control). To quantify the variability of the cycle and of the durations of the Up and Down states, we calculated their CV in the network. For the two networks (Fig. 6), the CV values were (typical (atypical)): 0.29 (0.34), 0.22 (0.25) and 0.20 (0.25) for the Down state, the Up state and the whole cycle duration, respectively, in the control condition; and 0.15 (0.15), 0.10 (0.20) and 0.13 (0.15) when GABA_B_ receptors were blocked. These values showed a substantial decrease in variability following GABA_B_ receptor blockade in both networks, especially for the duration of the Down states. Panels c and f in Figs. 6A and 6B illustrate the correlograms of the spike trains for the control and the GABA_B_blocked conditions.

**Figure 6A.**
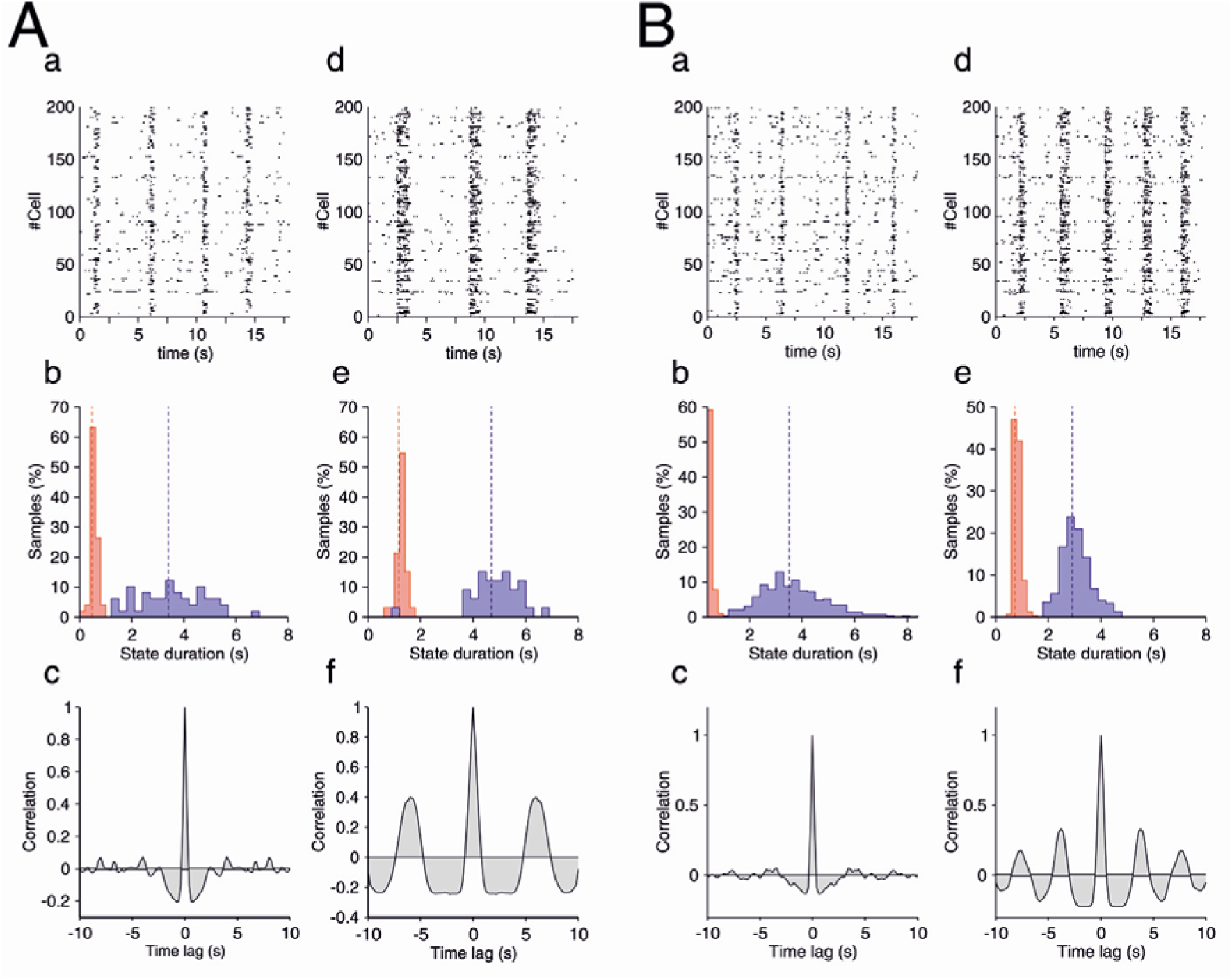
Properties of the slow oscillation in a typical (A) and atypical (B) networks. **A.** Plots on the left refer to the control condition and on the right to the slice with blocked GABA_B_ receptors. (a, d) Rastergrams. (b, e) Histograms of the duration of the Up (red) and Down (blue) states. Dashed lines correspond to their mean value: Mean durations of Up (Down) states are 0.45s (3.42s) in (b) and 1.17s (4.69s) in (e). (c, f) Spike-train correlation functions averaged over 100 pairs of neurons. Note how the oscillation becomes more regular in the blocked condition. **B.** Properties of the slow oscillation in an atypical network. Conventions are as in Figure 6A. Mean durations of Up (Down) states are 0.36s (3.51s) in (b) and 0.73s (2.91s) in (e).

### Explaining the modulation of variability by GABA_B_ receptors

In our experimental results we found that the decrease in the variability of the duration of Down states and complete cycles occurred in all cases, while the decrease in Up state duration variability only occurred in typical networks. To investigate the factors responsible for the variability of the Down state duration in our model, we looked at the traces of the synaptic and adaptation currents (Fig. 7A) for the two networks described in Fig 6.

**Figure 7.**
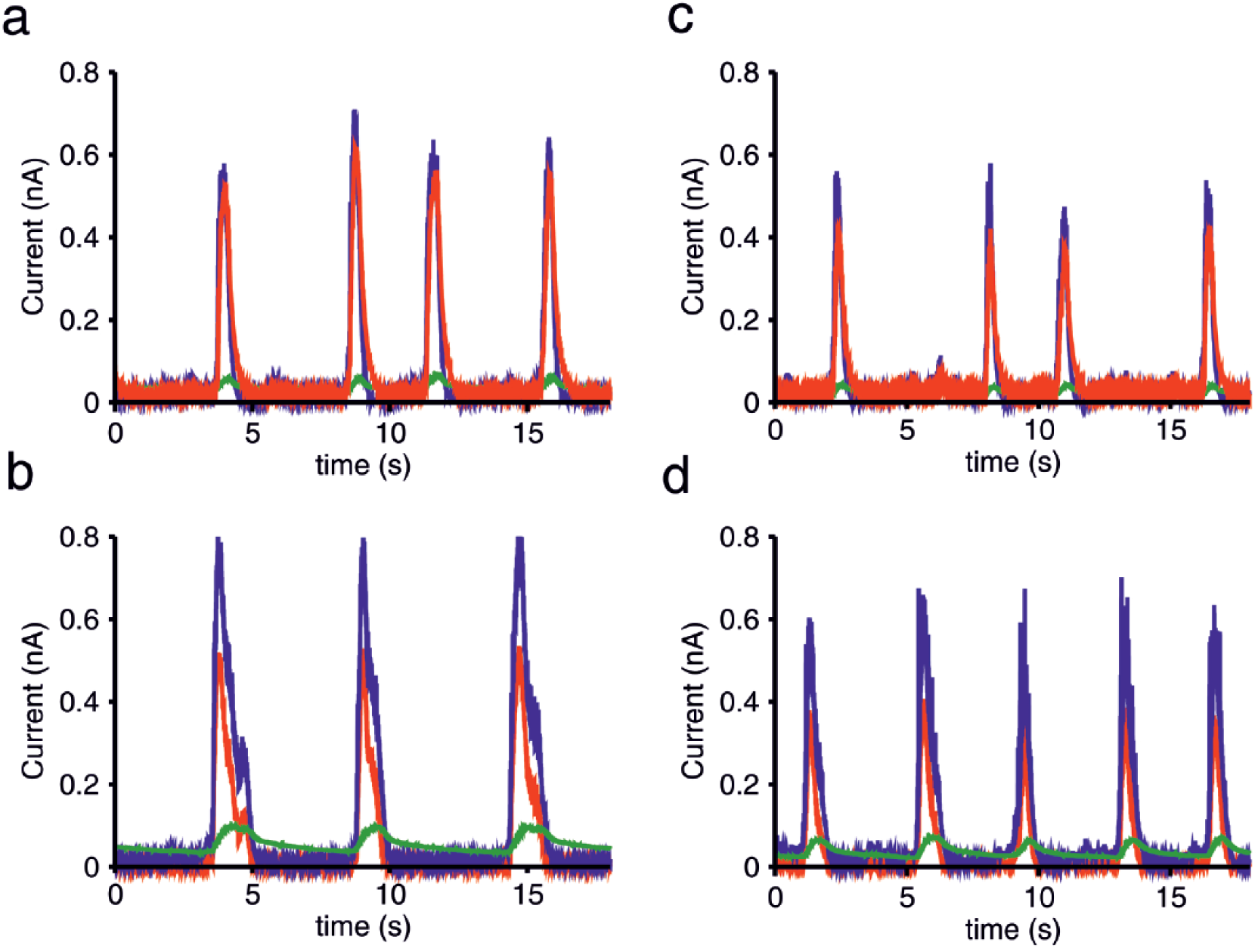
Temporal traces of the synaptic and adaptation currents. Temporal traces of the population averaged currents for the typical (left side) and the atypical (right side) network. Panels (a) and (c) refer to the control networks and panels (b) and (d) to the networks with the GABA_B_ receptors blocked. Notice the increasing regularity in the blocked case. Blue lines: total excitatory (AMPA plus NMDA) current. Red lines: total inhibitory (GABA_A_ plus GABA_B_) current. Green lines: adaptation current.

We first examined why SWO were rather regular when GABA_B_ receptors were blocked. Fluctuations of the neuronal excitation occur either from synaptic or external noise. Since in the GABA_B_-blocked condition the inhibition is fast (GABA_A_-mediated), fast inhibition can track these fluctuations easily (Compte et al., 2009; Renart et al., 2010) so that they do not propagate through the network unless a large population of excitatory neurons becomes active. Thus, the dynamics consist in a gradual increase of the excitation that starts during the Down state and grows until an Up state is generated (Fig. 7b and 7d). At this point the fast inhibition follows this large change in the excitation but cannot suppress it. During Up states, the adaptation current progressively increases and produces the end of the Up state. Since the characteristic time of the adaptation conductance is large and spiking is rare during the Down state, this conductance decays smoothly and slowly. These mechanisms give rise to a rather regular sequence of cycles (Fig. 7A).

When GABA_B_ receptors are not blocked, as in the case of the experimental control condition, these receptors produce two main effects. First, the total inhibitory current increases. A consequence of this increase is the shortening of the duration of the Up states (Fig. 6Aa,Ba and Fig. 7a,c). The second effect is the loss of regularity. Our experimental observations showed that when GABA_B_ receptors were not blocked, Down states could either be longer or shorter than in the blocked condition, the second case being the most typical. In both, the typical (Fig. 6A) and the atypical (Fig. 6B) simulated networks, the variability was higher in the control condition, but the origin of this variability has to be explained differently because the mean duration of their Down states was related differently to the corresponding networks with blocked GABA_B_ receptors.

For the atypical network (Fig. 6B), comparison of the temporal traces of the currents for the control and the blocked GABA_B_ networks obtained with simulations (Fig.7Ac and Ad) indicate that some Up states were suppressed in the control condition. This increased the mean duration of the Down states (Fig. 6B) and increased their variability (from CV=0.15 to 0.34). For the typical network (Fig. 6A) the temporal traces of the currents (Fig. 7a and b) indicate that, in contrast to what happened for the atypical case, now new Up states appeared with respect to the blocked condition. This also introduced a similar change in the variability of the duration of the Down states (CV increased from 0.15 to 0.29). To explain this different behavior in typical vs. atypical networks, let us focus on the way that the two networks were constructed. These two networks differ in the value of only two parameters: the NMDA conductance and the characteristic time of the adaptation conductance.

In the typical network, the NMDA unitary conductance is 40% larger than in the atypical one. To see the effect of a larger NMDA conductance on the duration of the Down states, let us consider a network identical to the atypical one (Fig. 6B) but with the NMDA unitary conductance increased by 40%. The stronger excitatory recurrent inputs reduced the duration of Down states in both, the control (Fig. 8A) and the blocked GABA_B_ (Fig. 8B) conditions. This effect can be observed by comparing the mean duration of Down states for the modified network with the corresponding mean duration for the original atypical network (Fig. 8A and B). However, one important difference arises: in the control condition, the shortening of the duration of the Down states is about 68% while in the blocked GABA_B_ condition it is only about 33%. Notice that this difference makes the original atypical network (Fig. 6B) become typical, in the sense that now the duration of Down states is longer when GABA_B_ receptors are blocked.

**Figure 8.**
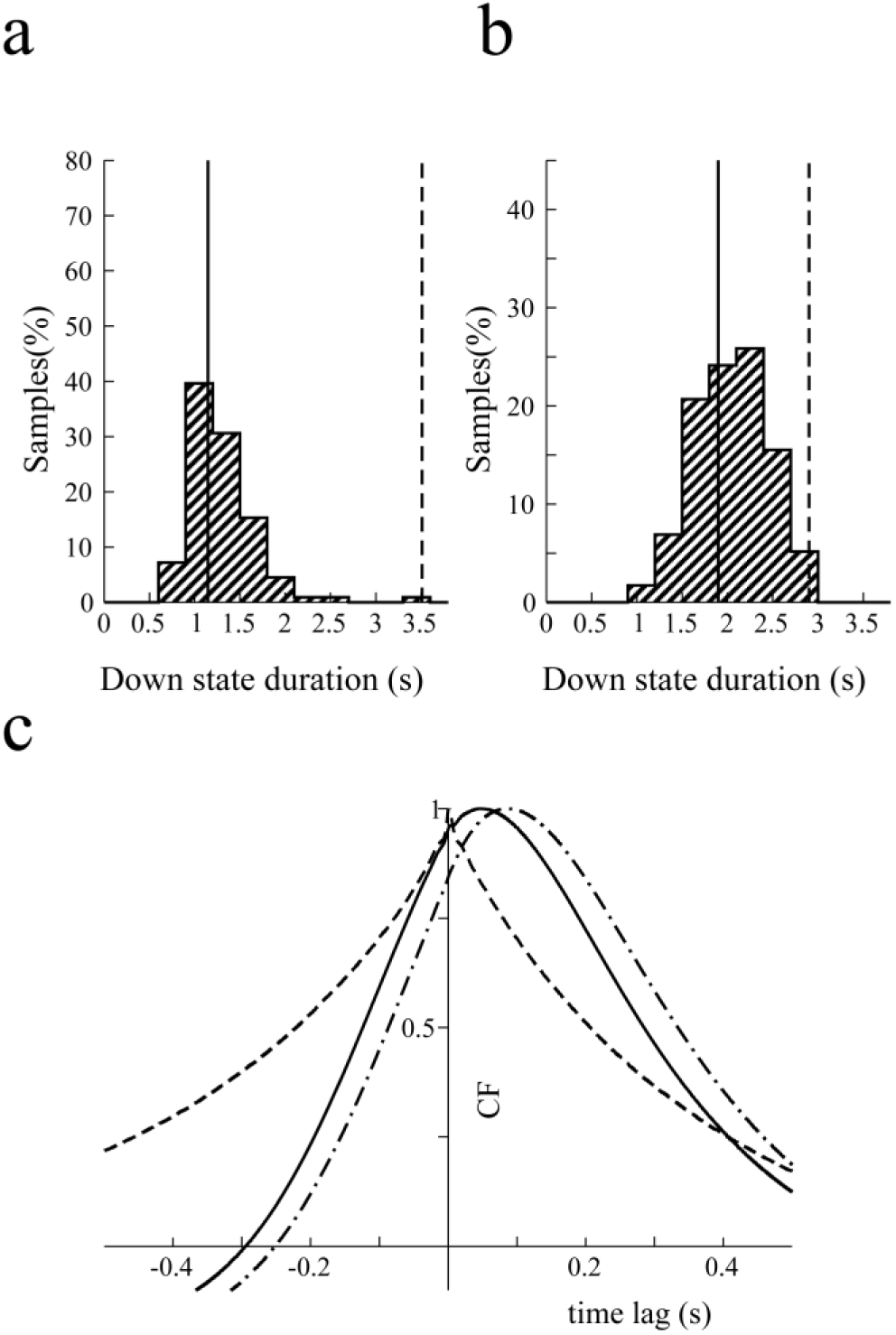
Effect of NMDA conductances on the duration of Down states. (a) Histograms of the duration of the Down states in the control condition of a network with the same parameters as the atypical one (Fig. 6B) but with NMDA unitary conductances increased 40%. Average Down state duration (solid line) is 1.1s, 68% smaller than in the original network, 3.51s (dashed line). (b) Same as (a) but in the blocked condition. Average Down states duration (solid line): 1.8s, about 33% smaller than in the original network, 2.91s (dashed line). (c) Correlation function between excitatory and inhibitory currents in the atypical network. Full line: Correlation function between total excitatory and inhibitory currents in the control condition. Dashed line: same as before but in the GABA_B_ blocked condition. Dashed-dotted line: Correlation function between total excitation and GABA_B_ component of the inhibition. CF, correlation function.

Why this differential shortening of Down states? The explanation can be found in the cross-correlation function between the excitatory and inhibitory currents in the atypical network (Fig. 8C). In the blocked GABA_B_ condition, the tracking of excitatory currents by inhibitory ones is almost instantaneous (the peak is located at 1.0 ms). A consequence of this is that inhibitory inputs can follow excitatory ones, until they are strong enough to cause the network to arrive to an Up state. However, in the control condition, the slow dynamics of the GABA_B_ receptors make the tracking of the excitation by inhibition difficult. This fact is reflected in the peak of the cross-correlation function at 46.0 ms (Fig. 8C, full line). This lag is 86.0 ms if only the GABA_B_ component of the inhibition is considered (dashed-dotted line). Since the tracking mechanism is not efficient in the control condition, the shortening of Down state duration is much more pronounced than in the blocked GABA_B_ condition.

The two differences between these two sample simulated networks were NMDA conductance, which we have already described, and the time of the adaptation conductance. Both networks (Fig. 6) were selected such that they had approximately the same Down state duration in the control condition. Although the network in Fig. 8A,B is already a typical network, the mean duration of its Down states in the control condition is shorter than that of the atypical network in Fig. 6B. To obtain the typical network in Fig 6A, we then made the characteristic time of the adaptation conductance 50% larger than in the atypical network.

In conclusion, the tracking of fast inhibitory currents to the excitation in cortical networks (Compte et al., 2009; Renart et al., 2010) is therefore spoiled by the presence of GABA_B_ currents spoiled this property.

## Discussion

In this paper we investigated the role of GABA_B_ receptors on the slow oscillatory rhythmicity driven by the alternating between Up and Down states. To that end we combined extracellular recordings of spontaneously active cortical network slices and computational experiments to further understand the mechanisms underlying slow wave activity. We found that GABA_B_ receptors controlled the duration of the active periods or Up states, such that their blockage elongated them, as previously described (Mann et al., 2009). We found that this effect was not mediated by the control of the firing rate during the Up state, but by contributing to the Up to Down transition, thus controlling the network synchronization. Furthermore, GABA_B_ receptors also had an impact on the subsequent silent periods or Down states, therefore modulating the complete oscillatory cycle. Interestingly, the effect of GABA_B_ receptor-blockade on the duration of the Down states can be elongation (most commonly), but also shortening. We explore in our computer model how these two opposing effects can be caused by the same intervention. The regularity of the oscillatory cycle is another parameter of the Up/Down dynamics that is modulated by GABA_B_ receptors, such that their blockade enhances the regularity and their activation introduces dynamical richness.

Although several biophysical mechanisms explaining SWO have been proposed, a full characterization of each one’s effects and a systematic study of their interactions is still missing. The difficulty with such studies is the fact that SWO is a spontaneous activity that emerges from the network and most mechanisms interact and influence globally the network dynamics, therefore the precise dissection of individual mechanisms is not an easy task. For example, a mechanism that only affects the firing rate during Up states, is going to modify not only Up but also Down states, since they are dynamically related. Further, the fact that one mechanism investigated by an external intervention (e.g. an agonist/antagonist) introduces a change in the Up/Down dynamics, does not guarantee the extent of its participation under physiological conditions or its interactions with other mechanisms. It is for this reason that these mechanisms are not yet well known and also why we need the use of computational models alongside the experiments to better explore a larger parameter space and mechanistic interactions.

The blockade of GABA_B_ inhibition resulted in this study and in others (Mann et al., 2009; Craig et al., 2013) in a consistent elongation of the Up states. Were GABA_B_ blockade to result in a decreased firing rate of the network, the Up state elongation could be seen as an indirect effect. However, the absence of effect on the firing rate points to a direct role of GABA_B_ inhibition in the termination of Up states. We observed that the average elongation (both in typical and atypical cases) of Up states was up to 182%, which in absolute terms is an elongation from an average of 0.56 (control) to 0.93 (after CGP 35348) seconds. Out of 37 cases, only 3 of them did not show an elongation of the Up state duration as a result of GABA_B_ blockade. All the rest elongated in a range that varied between 111% and 368%. Interestingly, this variability was independent of the original duration of Up states, which varied between 0.21 and 1.31 seconds, suggesting that GABA_B_ does not have a preferential role in Up state termination for either short or long Up states. Elongation was also independent of firing rate during Up states. This suggests that GABA_B_ is an independent mechanism that terminates Up states by acting in cooperation with other mechanisms, as it is the case for GABA_A_ receptors. The blockade of fast inhibition, mediated by GABA_A_ receptors results in increased firing rates in the Up states, that efficiently activate activity dependent mechanisms, such as potassium currents, that induce the termination of Up states, and shorten them (Sanchez-Vives et al., 2010). The role of GABA_A_ receptors on the termination of Up states and the initiation of the Down states suggests that they may as well have a role in the so-called Off-periods that disrupt local causal interactions in the cortical network in unresponsive wakefulness syndrome and natural sleep (Rosanova et al., 2018).

The impact of GABA_B_ inhibition on the variability of intervals also suggests that the participation of GABA_B_ in the termination of Up states varies depending on the functional state of the network, decreasing for those states that are highly regular. In the experimental study, we found that the regularity of the cycle significantly increased when GABA_B_ receptors were blocked. In the model, by blocking GABA_B_ the network went into an oscillatory alternation of states dominated by adaptation mechanisms, resulting in a more regular oscillatory rhythm. This shows that GABA_B_ activation not only plays a role in the termination of Up states, but also introduces variability in the oscillatory cycle. That the regularity of the oscillatory cycle can range from very high in deep (slow-wave) sleep or anesthesia to very low and chaotic during periods of transition to the awake state (Deco et al., 2009) suggests that different mechanisms regulate transitions between Up and Down states in different functional states. According to our results, a reduced contribution of GABA_B_ inhibition to the dynamics of the SWO would be expected in highly regular periods of activity such as deep sleep or anesthesia which, according to our model, could be well regulated by adaptation mechanisms. Experiments in awake rodents have shown different degrees of network synchronization: whereas alert states are characterized by desynchronized activity, resting awake states are characterized by more synchronized activity, with slow spontaneous fluctuations (Poulet and Petersen, 2008; Okun et al., 2010). GABA_B_ receptors could be involved in switching between these different functional states by modulating the network synchronization.

It has been shown that electrical stimulation of layer I is effective in terminating Up states (Mann et al., 2009). This effect is blocked by the GABA_B_ receptor-blocker CGP55845, suggesting that the Up state termination is mediated by GABA_B_ activation triggered by a subtype of interneuron in layer 1 called neurogliaform cell (Hestrin and Armstrong, 1996; Olah et al., 2007). However, our experimental results show that the termination of the spontaneous Up states seems to be independent of layer 1-mediated activation. Different roles for GABA_B1a_ and GABA_B1b_ subunits have also been proposed. GABA_B1a_ is preferentially located presynaptically and seems to be involved in spontaneous Up state termination; GABA_B1b_, on the other hand, is related to afferent or electrical stimulation via layer 1 activation (Craig et al., 2013). In our experiments, the disconnection of layer 1 from the slices did not result in the elongation of Up states; instead, Up state duration did not change or in some cases became shorter. However, applying GABA_B_ blocker CGP 35348 after removing layer 1, Up states increased their duration as it occurred in slices where layer 1 was not removed (Fig. S1). This result is in agreement with previous ones showing that spontaneous Up-to-Down transitions are independent of layer 1 activation (Craig et al., 2013). Craig et al. (2013) suggested that the change in the Up-to-Down transition slope is mediated by presynaptic GABA_B1a_ receptor activation. In this operational framework, our model predicted that changes in NMDA conductance together with firing rate adaptation are enough to generate activity in two different networks, similar to the ones we observed experimentally after blocking GABA_B_ receptors, as we showed for the typical and the atypical cases.

We also analyzed the effect of blockage of GABA_B_ in supragranular layers and the result was similar, showing that the modulation of the activity by GABA_B_ persisted in different layers and also in different cortical areas, in our case visual and prefrontal, and compatible with other authors’ and our own work (Mann et al., 2009; Wang et al., 2010; Craig et al., 2013).

We previously described that the partial blockade of GABA_A_ receptors with bicuculline increases the firing rate during Up states and decreases their regularity in active ferret cortical slices (Sanchez-Vives et al., 2010). More recently, Busche and colleagues showed, in wild-type anesthetized mice, that small concentrations of gabazine (a GABA_A_ receptor antagonist) desynchronize the network activity between distal cortical areas, and treatment with benzodiazepine (GABA_A_ agonist) restored the synchronization in a mouse model of Alzheimer’s disease characterized by low levels of synchronization in the control condition (Busche et al., 2015). Here, we show how the blockade of GABA_B_ receptors increases the regularity of the SWO, suggesting that GABA_B_ can introduce desynchronization in normal conditions. On this basis, we propose a model in which GABA can modulate the network synchronization by means of the activation of GABA_A_ and GABA_B_ receptors which generate opposing effects, synchronizing or desynchronizing the activity, respectively.

Activity-dependent adaptation, mediated by hyperpolarizing currents, has been proposed as a critical mechanism for the termination of Up states and maintenance of Down states. Such currents would be Ca^2+^ and Na^+^-dependent K^+^ currents (Compte et al., 2003; Sanchez-Vives et al., 2010) or AMPc-dependent potassium currents (Cunningham et al., 2006). Hyperpolarizing currents can also interact with other mechanisms such as synaptic depression, modulating the emerging patterns (Benita et al., 2012). Adaptation has also been considered in the dynamics of Up/Down states as a necessary mechanism, but in a more ample sense, such that it could include either hyperpolarizing ionic currents but also synaptic inhibition (Mattia and Sanchez-Vives, 2012). Here, in our model we considered firing rate adaptation and GABA_B_ receptors as the two dominant biophysical factors responsible for the termination of Down states and used experimental and modeling work to investigate how they participate in slow oscillations.

The model is a generalization of the standard leaky integrate-and-fire model in which a nonlinear current has been included (Parga and Abbott, 2007). Adaptation is taken as a linear firing adaptation current with a characteristic time appropriate for generating oscillations with an adequate frequency. A more complete way to describe adaptation in these slices is through an activity-dependent mechanism based on Na^+^ and Ca^2^-dependent K^+^ currents (Compte et al., 2003). In this case, spike firing during Up states induces the accumulation of Na^+^ and Ca^2+^ ions inside the axon, which in turn causes K^+^ ions to move outside the axon, hence hyperpolarizing the neuron and terminating Up states. The duration of this hyperpolarization is determined by the time course of the decay of the Na^+^ and Ca^2+^ concentrations (Wang et al., 2003), giving rise to Down states. However, a modeling framework where a simpler activity-dependent adaptation is responsible for the Up-to-Down state transitions produces, in the absence of GABA_B_ receptors, oscillations as regular as those observed in the slices. Our model considers firing rate adaptation and slow inhibition by GABA_B_ as the two biophysical elements determining the SWO; another plausible mechanism is short-term synaptic dynamics (Timofeev et al., 2000; Melamed et al., 2008; Benita et al., 2012); however, we did not need it to explain the slice behavior.

In conclusion, using *in vitro* experiments and computer models, we found that GABA_B_ receptors critically control the synchronization of the network discharge. According to our results the decrease in GABA_B_ receptor activation enhances the cycle regularity and Up state duration. This suggests that in normal conditions, GABA_B_ is a source of desynchronization in cortical activity.

## Acknowledgements

This work was supported by EU H2020 Research and Innovation Programme, Grant 785907 (HBP SGA2) and BFU2017-85048-R (MINECO). We would like to thank Tony Donegan for editing the paper.

---

**Figure S1.**
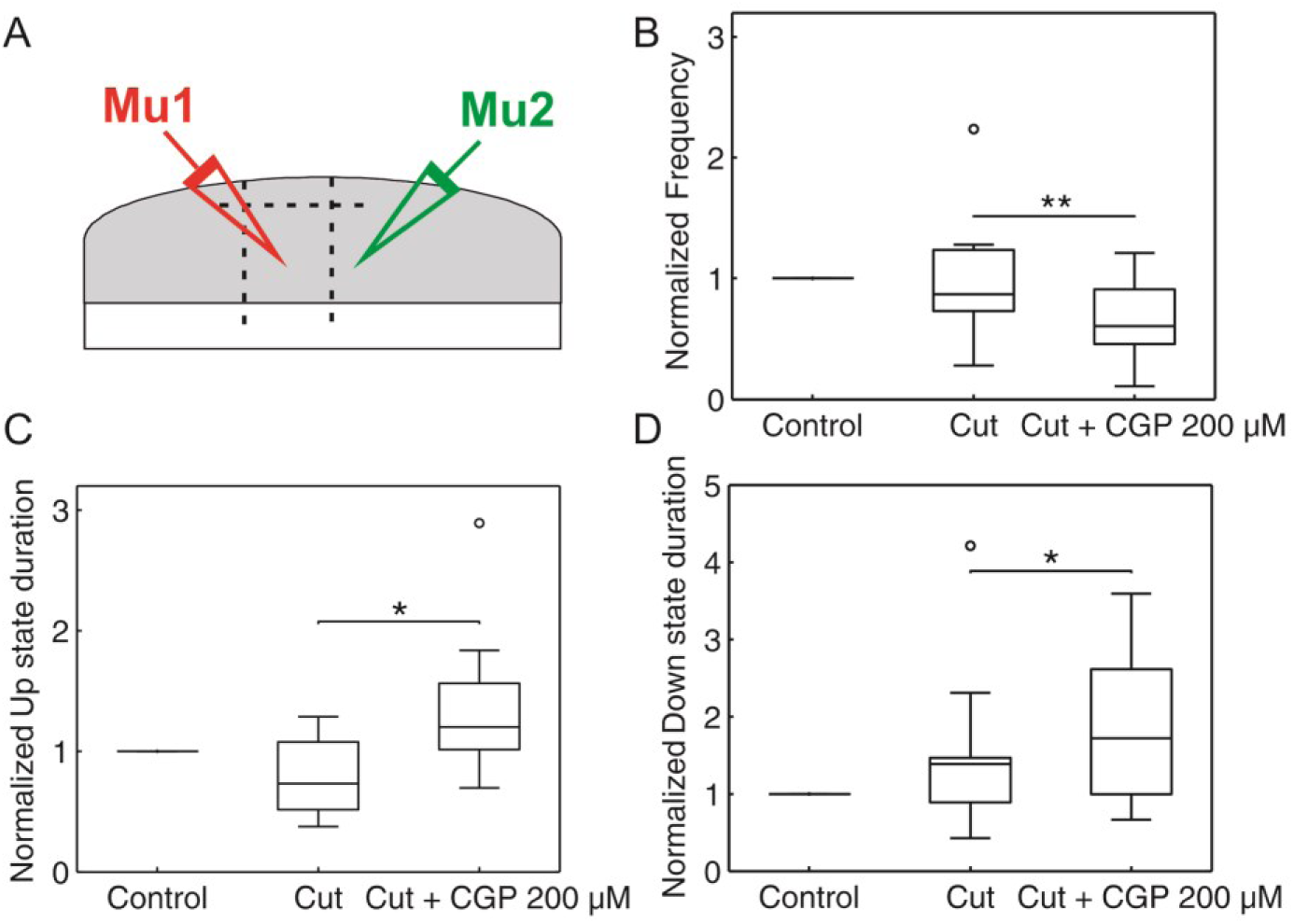
Effects of elimination of layer 1. A. Experimental sketch with double extracellular recordings. B–D. Average of normalized frequency (B), Up state duration (C), and Down state duration (D) in control, after layer 1 cut, and after application of CGP 200µM. A Student’s *t*-test was used to evaluate how different the normalized averages were from 1.0., *p* < 0.05 (*); *p* < 0.01 (**); *p* < 0.001 (***). (*n*= 13).

